# Differential patterns of change in brain connectivity resulting from severe traumatic brain injury

**DOI:** 10.1101/2021.10.27.466136

**Authors:** Johan Nakuci, Matthew McGuire, Ferdinand Schweser, David Poulsen, Sarah F. Muldoon

**Affiliations:** Neuroscience Program, University at Buffalo, Buffalo, NY; Department of Neurosurgery, University at Buffalo, Buffalo, NY; Buffalo Neuroimaging Analysis Center, Department of Neurology, School of Medicine and Biomedical Sciences, University at Buffalo, Buffalo, NY; Center for Biomedical Imaging, Clinical and Translational Science Institute, University at Buffalo, Buffalo, NY; Department of Mathematics and CDSE Program, University at Buffalo, Buffalo, NY

**Keywords:** traumatic brain injury, structural connectivity, network theory, post-traumatic epilepsy

## Abstract

**Background:** Traumatic brain injury (TBI) damages white matter tracts, disrupting brain network structure and communication. There exists a wide heterogeneity in the pattern of structural damage associated with injury, as well as a large heterogeneity in behavioral outcomes. However, little is known about the relationship between changes in network connectivity and clinical outcomes.

**Methods:** We utilize the rat lateral fluid-percussion injury (FPI) model of severe TBI to study differences in brain connectivity in 8 animals that received the insult and 11 animals that received only a craniectomy. Diffusion Tensor Imaging (DTI) is performed 5 weeks after the injury and network theory is used to investigate changes in white matter connectivity.

**Results:** We find that 1) global network measures are not able to distinguish between healthy and injured animals; 2) injury induced alterations predominantly exist in a subset of connections (subnetworks) distributed throughout the brain; and 3) injured animals can be divided into subgroups based on changes in network motifs – measures of local structural connectivity. Additionally, alterations in predicted functional connectivity indicate that the subgroups have different propensities to synchronize brain activity, which could relate to the heterogeneity of clinical outcomes.

**Discussion:** These results suggest that network measures can be used to quantify progressive changes in brain connectivity due to injury and differentiate among subpopulations with similar injuries but different pathological trajectories.

**Impact Statement:** White matter tracts are important for efficient communication between brain regions and their connectivity pattern underlies proper brain function. Traumatic brain injury (TBI) damages white matter tracts and changes brain connectivity, but how specific changes relate to differences in clinical/behavioral outcomes is not known. Using network theory to study injury related changes in structural connectivity, we find that local measures of network structure can identify subgroups of injured rats with different types of changes in brain structure. Our results suggest that these different patterns of change could relate to differences in clinical outcomes.

## Introduction

White matter tracts provide the underlying structure for communication between brain regions and therefore damage to these tracts can have significant impact on brain function. Traumatic Brain Injury (TBI) can cause diffuse axonal injury beyond the site of the trauma, with white matter tracts particularly vulnerable to injury (Johnson et al., 2013b). The location and extent of an injury can be assessed with diffusion Magnetic Resonance Imaging (dMRI) and alterations in diffusion properties can characterize structural changes in the brain (Basser et al., 1994).

In humans (Benson et al., 2007; Kraus et al., 2007; Mac Donald et al., 2007; Wilde et al., 2008; Kinnunen et al., 2011) and rodents (Mac Donald et al., 2007; Budde et al., 2011; van de Looij et al., 2012; Pischiutta et al., 2018), the diffusion metrics of fractional anisotropy (FA), axial diffusivity (AD), radial diffusivity (RD), and mean diffusivity (MD) have all shown injury related changes in brain structure and additional changes have been found in white matter fiber density (Harris et al., 2016a; Wright et al., 2017). In animal models, the changes in diffusion metrics and fiber density indicate the possibility that, alongside axonal and myelin degeneration, there is axonal sprouting and structural reorganization following an injury (Christman et al., 1997; Harris et al., 2016a). Despite the extensive research in TBI induced structural changes, thus far it has been difficult to link observed changes in brain structure to behavioral/clinical outcomes because the extent of damage and change in brain structure is highly heterogeneous in humans and rodents.

Diffusion metrics can provide information about injury related changes to brain structure, and the use of tools from network theory (Bassett and Sporns, 2017) offers an opportunity to understand how TBI induces local and global changes in brain connectivity (Feldt et al., 2011; Sharp et al., 2014; Bassett and Sporns, 2017). Studies utilizing structural networks have found that TBI induces changes in a subset of brain connections (Hayes et al., 2016; Iraji et al., 2016; Thengone et al., 2016; Dall’Acqua et al., 2017) and can alter a network’s characteristic path length, global/local efficiency, betweeness centrality, eigenvector centrality, and small-worldness (Caeyenberghs et al., 2014; Yuan et al., 2015; Königs et al., 2017; van der Horn et al., 2017). Furthermore, it has been shown that betweenness- and eigenvector-centrality can potentially be used as diagnostic biomarkers (Fagerholm et al., 2015).

Importantly, in addition to the observed heterogeneity of changes in network structure, there is also a large heterogeneity in clinical outcomes post-TBI, ranging from cognitive impairment (Kinnunen et al., 2011) to an increased risk of developing post-traumatic epilepsy (PTE) (Annegers et al., 1998). The incidence of PTE can be as high as 53% of patients, depending on injury severity and type of injury (penetrating vs blunt) (Frey, 2003), and correlates with the severity of the injury (Annegers et al., 1998; Christensen et al., 2009; Mahler et al., 2015). Unfortunately, there are no clear biomarkers for predicting clinical outcomes post-TBI (Pitkänen and Immonen, 2014).

Here, we use the lateral fluid-percussion injury (FPI) model to induce severe TBI in rats and use network theory to study the associated changes in brain structure post-injury (McIntosh et al., 1989; Rau et al., 2012, 2014; Smith et al., 2018). This model mimics the damage observed clinically (Xiong et al., 2013) and displays a heterogeneity in the severity of injury both structurally and functionally. In rats, FPI induced neuronal injury is marked by a combination of focal cortical contusion and diffuse subcortical neuronal injury (Hicks et al., 1996). Additionally, the FPI model produces behavioral and cognitive deficits, such as movement and memory deficits, that are commonly seen in patients with TBI (Hamm, 2001; Morales et al., 2005), and some rats will go on to display spontaneous seizures (Bolkvadze and Pitkänen, 2012; Immonen et al., 2013).

To characterize the heterogeneity in structural damage post-injury, we examine the properties of large-scale structural brain networks derived from dMRI imaging data in injured and control rats. We find differences in subnetwork brain structure between injured and control rats and that animals within the TBI, but not control, group can be clustered into two distinct subpopulations driven by changes in local motif structure of the brain networks. Further, simulated functional connectivity patterns predict that one subgroup shows higher levels of synchronization, which we speculate could be related to differences in clinical outcomes between subgroups such as the propensity to later develop epilepsy.

## Materials and Methods

### Ethical Treatment of Animals

The Institutional Animal Care and Use Committee at SUNY Buffalo approved all procedures in these studies.

### TBI Procedure

Adult, male, Wistar rats (300-350g) were obtained from Charles River Laboratories (Wilmington, MA) and single housed in a 12-hour light/dark schedule with ad libitum access to food and water. The lateral fluid percussion injury procedure was performed as we have previously published (McGuire et al., 2019).

Briefly, animals were deeply anesthetized using 2–4% isoflurane. A 5 mm trephin was used to make a craniectomy over the right hemisphere equidistant between the lambda and the bregma and adjacent to the lateral ridge. Animals received a 20 ms fluid pressure pulse to the dura at an average of 2.79ATMs of atmospheres (range of 2.64 to 2.98 ATMs). Following injury, animals became apneic after injury for an average of 38 seconds (range of 0 to 60 seconds) and were manually ventilated with supplemented O_2_ until normal breathing resumed. A death rate of 33% was observed for injured animals. Righting reflex time was recorded for each animal, averaging 34 minutes (range of 17 to 60 minutes). Injury severity was determined based on functional behavior as measured by the neurological severity score (NSS), assessed at 24 hours after injury. In total 8 animals received the insult (TBI Group) and 11 animals received only the craniectomy (Control Group). Given the small sample size, the severe model of FPI was utilized to maximize the heterogeneity of injury in the TBI group.

### Diffusion Tensor Imaging

To record changes in brain network structure resulting from TBI, Diffusion Tensor Imaging (DTI) scans were performed 5 weeks after the insult with a 20 cm diameter horizontal-bore 9.4 Tesla magnet (Biospec 94/20 USR, Bruker Biospin) equipped with imaging gradient coils supporting 440 mT/m gradient strength and 3440 T/m/s maximum linear slew rate (BGA-12S HP; Bruker Biospin). We employed a cross-coil configuration with a 4-channel receiver rat-brain surface array and a transmitter volume coil. The scanner was operated with ParaVision (version 5.1; Bruker Biospin). The DTI scans consisted of a respiration-gated 3D Spin-Echo DTI sequence with a two-shot echo-planar readout and reversed readout gradients for Nyquist ghost suppression. DTI acquisition parameters were as follows: 213×144×166 μm^3^ field of view; 133×90×52 matrix zero-filled to 266×180×104; 1 average; TE/TR 25.96ms/1000ms; B=1000 s/mm2 in 30 directions across the whole sphere; 4 unweighted acquisitions (B0); 3ms diffusion times at 415.9 mT/m and 12ms duration. The acquisition time with gating was approximately 90 to 120 minutes.

### DTI Preprocessing and Structural Connectivity

Voxel dimensions of the diffusion images were scaled by a factor of 10x. This scaling factor is only applied to the header information in the DTI file in order to make the dimensions of the voxels compatible with the FSL software package which expects the DTI images to have human sized voxels. It is important to note that scaling does not change the underlying DTI images.

Motion and eddy-currents were corrected with FSL Eddy (Andersson and Sotiropoulos, 2016). Post eddy and motion correction, B0 images were averaged to create a brain mask using FSL Brain Extraction Tool (BET) (Smith, 2002). Advanced Normalization Tools (ANTs) (Avants et al., 2008), specifically ANTs Syn with default parameters, was used to register a brain atlas with 150 regions spanning cortical and subcortical areas (Valdés-Hernández, 2011) to the average of the B0s.

### Fiber Tracking and Network Construction

Fiber tracking was done in DSI Studio with a modified FACT algorithm (Yeh et al., 2013). Whole brain seeding was used to generate 500,000 tracts with a fractional anisotropy threshold of 0.12 and angular threshold of 45°, step size of 1mm, minimum length of 2mm, maximum length of 300mm based on the scaled voxel dimensions. In order to fully sample the fiber orientations within a voxel, tracking was repeated 1,000x with initiation at a random sub-voxel position generating 1,000 connectivity matrices per animal.

For each animal, an undirected weighted structural connectivity matrix, A, containing 150 brain regions was constructed by counting the number of streamlines between brain regions (Valdés-Hernández, 2011). The final connectivity matrix based on the average of the 1000 matrices. Connectivity matrices were then normalized by dividing the number of streamlines (*T*) between region *i* and *j*, by the combined volumes (*v*) of region *i* and *j*. This normalization is important to ensure that the strength of the connectivity between two regions is not influenced by volumes of the two respective regions (Zalesky et al., 2010b).

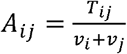

### Network Similarity

We assessed global dis-similarity between two subjects, *p* and *q*, based on the similarity of their structural connectivity using the L2-Norm which is defined as,

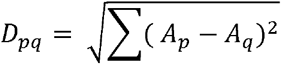

where *A*_*p*_ and *A*_*q*_ correspond to the structural connectivity matrices from subject *p* and *q*, respectively.

### Eigenvalue Spectrum

Each connectivity matrix (*A*) was decomposed to corresponding eigenvalues and eigenvectors. Eigendecomposition is obtained by solving the linear equation,

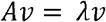

where λ = [ λ_1_ … λ_n_] are the corresponding eigenvalues (EigVal) and *v* = [*v*_*1*_*… v*_*n*_] the eigenvectors of *A*. The eigenvalue spectrum of *A* describes structural properties of the network.

Statistical differences were based on the full distribution of eigenvalues, pooled across TBI and Control animals respectively, and assessed with the Kolmogorov-Smirnov (K-S) test. We report the p-value and K-S test statistic (D) with a range [0, 1] which quantifies the maximum differences between the cumulative distributions.

### Network Based Statistics

Network-Based Statistics (NBS) was used to identify subnetworks which increased or decreased in strength due to trauma (Zalesky et al., 2010). Previous research has identified subnetworks that are significantly altered in moderate to severe TBI in humans (Dall’Acqua et al., 2017; Solmaz et al., 2017). We used NBS to test if similar changes occur in the FPI model of severe TBI in rats.

NBS implementation involves initially calculating the T-statistic for each connection between animals in the TBI and Control groups followed by applying a primary threshold to the T-statistic to identify a subnetwork. The threshold results in subnetwork with *L* edges. Statistical significance of the subnetwork is based on randomizing the group labels between individuals and calculating the size of the subnetwork after randomization. Familywise error (FWE)-corrected *p* values were calculated on the subnetwork using a null distribution derived from 5,000 permutations (Zalesky et al., 2010a). Subnetworks that survived a network-level threshold of p < 0.05 (FWE corrected) were considered significant. NBS analysis was conducted using the Brain Connectivity Toolbox (Rubinov and Sporns, 2010).

Since the choice of a primary T-statistic threshold is arbitrary, it is important to ensure the results are consistent over a range of thresholds. Therefore, we conducted our analysis with threshold values ranging from 1.0 to 2.5 in steps of 0.1. In the manuscript, we present the results with the strongest FWE-corrected p-value which occurred with a T-threshold of 1.8 for the subnetwork that increased in strength in the TBI group, and T-threshold of 2.0 for the subnetwork that decreased in strength in TBI group.

### Minimum Spanning Tree

The Minimum Spanning Tree (MST) is a subnetwork that connects all nodes while minimizing edge weights without forming loops (Kruskal, 1957; Tewarie et al., 2015). Since in brain networks, edge weights represent the density of tracts between regions and we are interested in information flow along these tracts, we calculated the MST using the inverse of the connections in structural connectivity. We additionally calculated the cost network associated with the MST (MST-Cost). Network cost is defined as the connection weight multiplied by the Euclidian distance between the centroids of two regions (Achard et al., 2006; Heuvel et al., 2012).

Statistical differences were based on the distribution of connections assessed with the Kolmogorov-Smirnov (K-S) test between the pooled TBI and Controls. We report the p-value and K-S test statistic (D) with a range [0, 1] which quantifies the maximum differences between the cumulative distributions.

### Degree

The weighted node degree (*k*_*i*_) is defined as the sum of all connections of a node (Rubinov and Sporns, 2010),

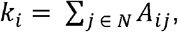

where *A* is the weighted adjacency matrix of a network with *N* nodes (Rubinov and Sporns, 2010).

### Betweenness-Centrality

Betweenness-Centrality of a brain region is the fraction of all shortest paths in the network pass through that region and is defined as,

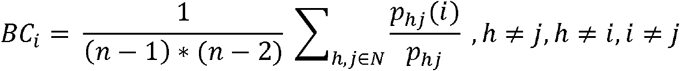

Where *p*_*hj*_ is the number of shortest paths between *h* and *j*, and *p*_*hj*_ is the number of shortest paths between *h* and *j* that pass through *i* (Rubinov and Sporns, 2010).

### Local Efficiency

Local efficiency (*E*_*l*_) is the global efficiency computed on the neighborhood of node *i*,

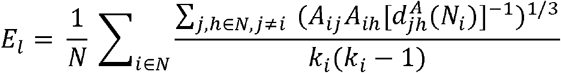

where *A*_*ij*_ and *A*_*ih*_ is strength of the connection between node *i* to *j* and *h*, respectively, and *d*_*jh*_ *(Ni)* is the length of the shortest path between node *j* and *h* that contains only neighbors of node *i* (Rubinov and Sporns, 2010).

### Motif Analysis

Motifs (*M*) are patterns of network connections among a specified number of nodes forming a subgraph (*g*), which act as building blocks for complex networks (Sporns and Kötter, 2004). Given that our structural connectivity matrices are undirected, our analysis focused on two types of three-node motifs and six types of four-node motifs. Individual motifs are named using the convention from Sporns and Kötter, 2004.

To assess how strongly a brain region participates in a specific motif, we calculated the motif coherence (*Q*) for each brain region (Onnela et al., 2005). We used motif coherence because it accounts for low probability values due to one connection weight being low versus all connection weights being low (Onnela et al., 2005). For each individual motif pattern, *g, Q* is defined as

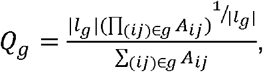

Where *l*_*g*_ is the set of edges in the specified subgraph, |*l*_*g*_| is the number of edges in that subgraph, and *A* is the connection strength between brain region *i* and *j*. Motif coherence was calculated using the Brain Connectivity Toolbox (Rubinov and Sporns, 2010).

### Z-score calculations

To assess differences in regional measures of brain structure (degree, betweenness-centrality, local efficiency, and motifs), we used a permutation analysis to calculate z-scores based on the difference in the TBI or T1/T2 subgroups when compared to the Control group. Specifically, for a single measure, *Q*, the difference between the TBI/T1/T2 and Control groups is

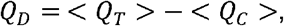

where <*Q*_*T*_> and <*Q*_*C*_> are the average of the measure across the TBI/T1/T2 and Control groups, respectively. Z-scores were then calculated by permuting group or subgroup labels 10,000x, and, *Z*_*Q*_ was calculated as,

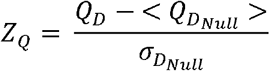

where 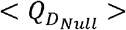 and 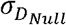 represent the average and standard deviation of the coherence for a motif subtype across the 10,000 permutations, respectively. The use of 10,000 permutations ensured that the tails of the null distribution were adequately sampled. Brain regions with Z-scores (*Z*_*Q*_) greater or lower than 1.96 (corresponding to p-values of 0.05 or lower) were considered statistically significant. When visualizing the regional motif coherence, we average the Z-scores of a node across all statically significant motif subtypes.

### Clustering Analysis

To test for the existence of subpopulations, we designed a two-step unsupervised clustering procedure that utilized network measures to cluster animals into subgroups. The clustering analysis was performed separately for TBI and Control groups. In the first step, using *kmeans*.*m* in MATLAB, we clustered animals based on similarity for each of the 16 network measures studied: Structural Connectivity (Wei), Eigenvalue spectrum (EigVal), NBS, MST, MST-Cost, Degree (Deg), Betweenness-Centrality (BC), Local Efficiency (Elocal) and 8 distinct Motifs. Similarity between animals was calculated using the L1-Norm. The optimal number of clusters (*K*) was determined by iterating *K* from 1 to N (N_TBI_ = 8; N_Control_ = 11, respectively) and the *K* with the maximum average silhouette value, which measures how similar a subject is to other subjects in the same cluster, was chosen as the optimal number of clusters.

To implement this, we first used only the structural connectivity of each animal to perform clustering, thus the animals were assigned to a cluster based only on the similarity in their structural connectivity. This was then repeated for the other network measures. The animals therefore were assigned a cluster label for each of the 16 network measures.

In the second step, we created, an association matrix that records the number of times two animals were in the same cluster across the 16 network measures. Subgroups within a population were identified by clustering the association matrix using modularity-maximization. Modularity-maximization identifies communities in which the similarity within a cluster is greater than between clusters. In addition, modularity-maximization does not require a predefined number of clusters to be specified. The resolution of the clusters can be controlled with parameter, γ.

Modularity-maximization was implemented with the Generalized Louvain algorithm (γ = 1) (Mucha et al., 2010). In order to avoid such suboptimal results due to initial seeding, clustering was repeated 100x and final results were based on the consensus across the 100 runs (Bassett et al., 2013).

### Predicted Functional Connectivity

Functional connectivity was predicted using the path transitivity (Goni et al., 2014). Path transitivity estimates the amount of signal that propagates from brain region *i* to *j*, while taking into account the number or re-entrants connections along the shortest path (Avena-Koenigsberger et al., 2017). Path Transitivity was calculated using the Brain Connectivity Toolbox (Rubinov and Sporns, 2010).

To assess changes in predicted functional connectivity, statistical significance was based on the difference in the TBI and subgroups when compared to the Control group. We first calculated the average predicted functional connectivity, *<FC*_*T/S*_*>*, for the TBI and for the subgroups followed by subtracting the average predicted functional connectivity from the Control, *<FC*_*C*_*>*.

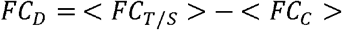

The resulting values represent the overall change in predicted functional connectivity *FC*_*D*_, for the TBI and subgroups, respectively. We then assessed differences between the TBI and the subgroups with an Analysis of Variance (ANOVA).

However, because this ANOVA does not assess which specific connections have a higher propensity to change, we performed additional analysis. To assess changes in individual connections, statistical significance was based on permuting group labels 10,000x and connections with Z-scores (*Z*_*FC*_) greater or lower than 1.96 from the null distribution (corresponding to p-values of 0.05 or lower) were considered statistically significant. Specifically, for each connection, *Z*_*FC*_ was calculated as

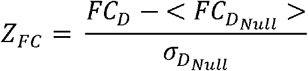

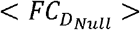 and 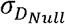 represent the average and standard deviation of the predicted functional connectivity across the 10,000 permutations, respectively.

### Experimental design and statistical analysis

The lateral fluid percussion model of TBI was used to induce a severe injury in Wistar rats (n = 8) and sham controls (n = 11) (Figure 1A). The sham controls received the surgery, but no injury. Five weeks after the injury, animals underwent DTI imaging (Figure 1B) from which tractography and structural connectivity was determined (Figure 1C-D).

**Figure 1.**
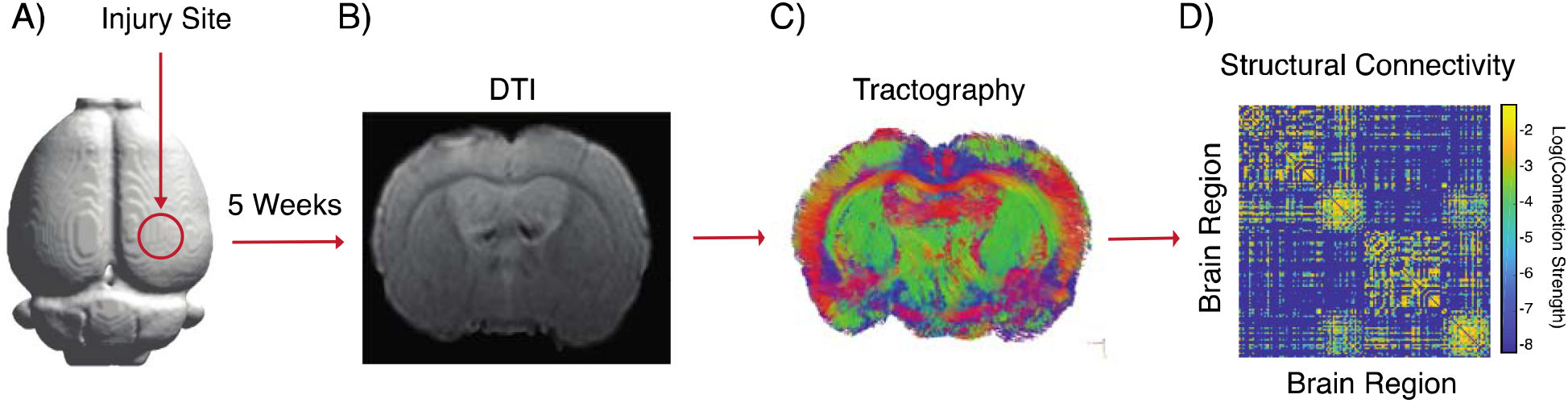
Experimental design and analysis. A) Injury was induced in the right hemisphere using the lateral fluid percussion model. B) Five weeks after injury animals underwent DTI imaging from which (C) white matter tracts and (D) structural connectivity was derived.

Network measures were calculated using the Brain Connectivity Toolbox (Rubinov and Sporns, 2010) and unique scripts in MATLAB 2017B (MathWorks). Statistical analyses were based on a combination of parametric and nonparametric tests as appropriate, and permutation testing was used when analyzing measures of local network structure in which multiple comparison issues applied. See individual methods sections for details. In addition, BrainNet Viewer was used to display results (Xia et al., 2013).

## Results

Here we quantify changes in structural connectivity in a population of 8 animals that received a severe-TBI using the lateral-fluid percussion injury (FPI) model and 11 control animals that received only the craniectomy surgery. Five weeks after the injury, animals underwent DTI imaging from which structural brain networks were created. We first compare brain network structure between control and TBI animals using network measures that assess network structure at (i) the global level (ii) subnetwork level and (iii) local level, and then show how these measures can be further used to detect the heterogeneity of injuries within TBI animals.

### Global Network Analysis

We first measured brain network structure at the global level in both TBI and Control animals in an attempt to identify types of large-scale structural changes associated with TBI. Specifically, we used the Euclidean distance between structural connectivity matrices to measure the magnitude of dis-similarity between the TBI and the Control group (Supplemental Figure 1A) but found no significant differences (Supplemental Figure 1B). We also examined the eigenvalue spectrum of the networks but again observed no differences between the TBI and Control groups (Supplemental Figure 1C). These results indicate that at the global level, these two measures of network structure are not able to distinguish between the two groups.

**Supplemental Figure 1.**
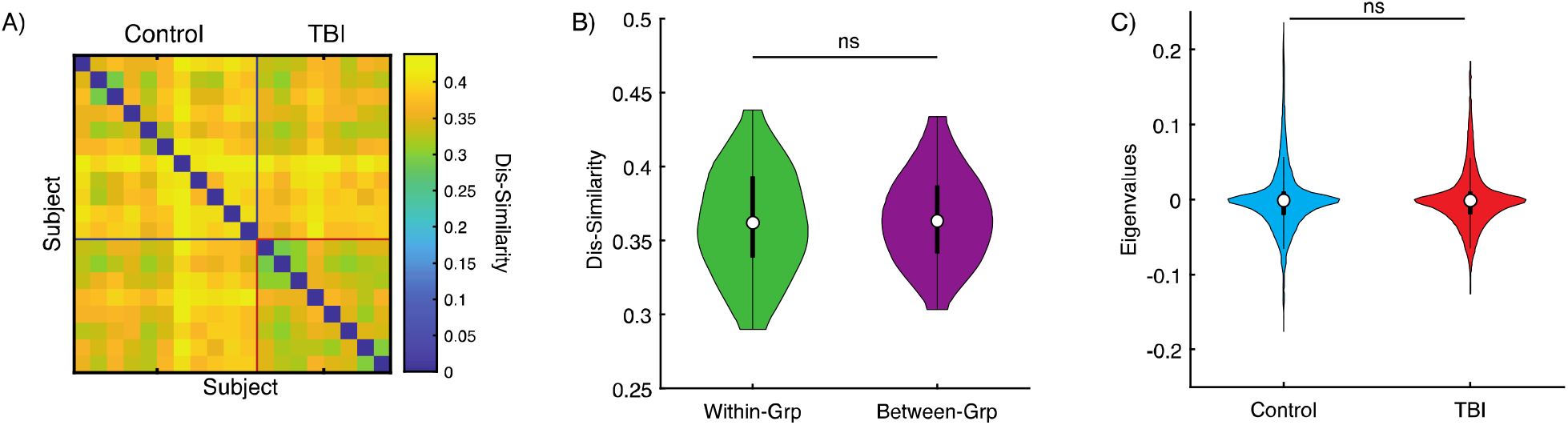
Global dis-similarity between groups. A) the magnitude of dis-similarity between all animals calculated using the Euclidian distance between structural connectivity networks. B) We observed no significant differences in the magnitude of Within- and Between-Group dis-similarity (T_169_ = -0.73, p = 0.46; Within-Grp: 0.36 ± 0.035 (SD); Between-Grp: 0.37 ± 0.029 (SD)). C) The distributions of Eigenvalue spectrums for TBI and Control groups were not different (Kolmogorov-Smirnov test; D = 0.016; p = 0.99). For each violin plot, the central mark indicates the median, the bottom and top edges of the box indicate the 25th and 75th percentiles, respectively and whiskers extend to the most extreme data points not considered outliers.

### Subnetwork Analysis

We next asked if the injury induced by the FPI primarily effects only a subset of connections in the brain. We first applied Network Based Statistics (NBS) to identify subnetworks that increased or decreased in strength in the TBI group relative to the Control group. With NBS, we identified a subnetwork consisting of 268 connections (n = 8 rats), which increased in strength (Figure 2A) (p << 0.001). In addition, the magnitude of the increase in strength of the connections was negatively correlated with the distance between brain regions (r = -0.46, p = 1.53 × 10^−15^) (Figure 2B). Alongside this component, NBS identified a smaller subnetwork with 74 connections (n = 8 rats) in which the connection strengths decreased (Figure 3C) (p << 0.001).

**Figure 2.**
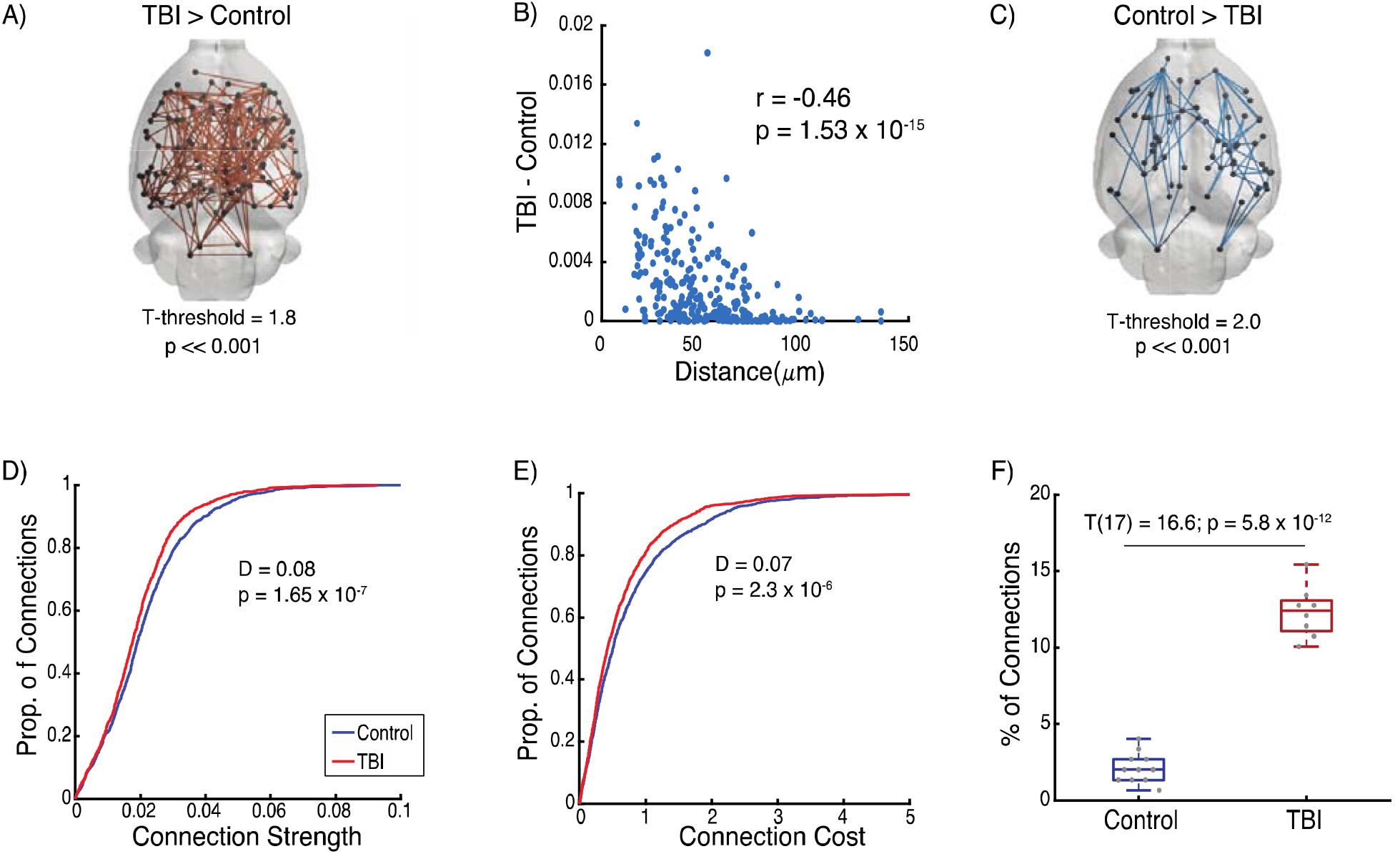
Injury induced changes in structural connectivity subnetworks. A) NBS identified a subnetwork increasing in connectivity strength in the TBI group. B) Correlation of magnitude differences of the subnetwork from (A) with distance between brain regions (r = -0.46, p = 1.53 × 10^−15^). C) Subnetwork of connections found with NBS that decreased in strengths in the TBI group. D and E) Differences in the proportion of connections in the TBI and Controls for the MST- (D = 0.08, p = 1.65 × 10^−7^) and MST-Cost subnetworks (D = 0.07, p = 2.3 × 10^−6^), respectively. F) Percentage of connections in the MST that are present in the NBS subnetwork for TBI and Controls (T_17_ = 16.6, p = 5.8 × 10^−12^). For each boxplot, the central mark indicates the median, the bottom and top edges of the box indicate the 25th and 75th percentiles, respectively and whiskers extend to the most extreme data points not considered outliers.

**Figure 3.**
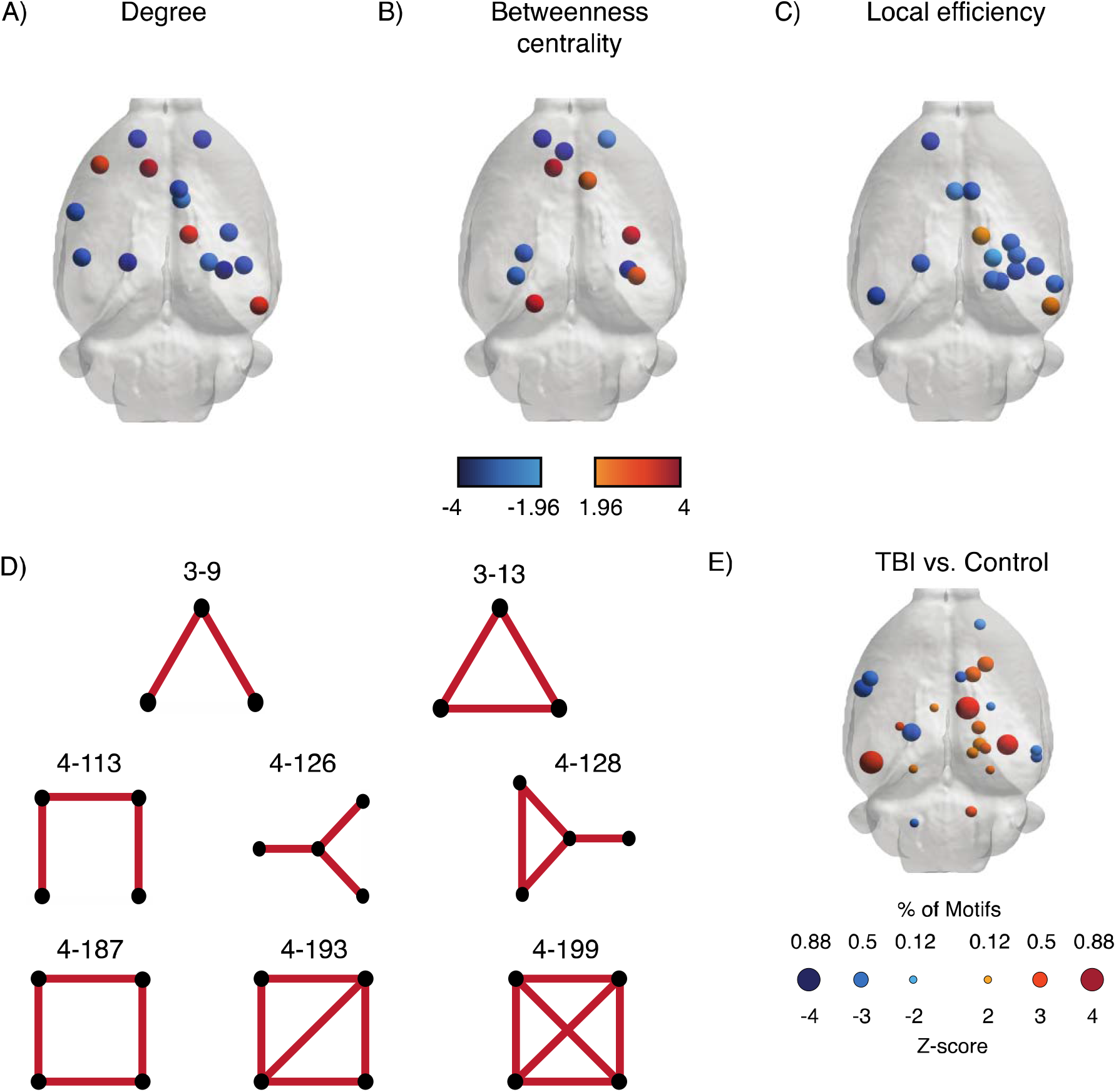
Local analysis of injury induced changes. Significant differences between TBI and Control groups in a brain region’s A) degree, B) betweenness-centrality, and C) local-efficiency. D) Schematic of 3- and 4-node motifs. E) Regions for which differences in motif coherence are statistically significant in the TBI group. Statistical significance was based on permutation testing with 10,000 randomizations. For (E), color corresponds to the average Z-score of all significant motifs and size corresponds to percentage of motifs which a brain region was significantly different. Only brain regions with |Z-score| > 1.96 corresponding to a p-value < 0.05 are shown.

We also assessed if there were corresponding changes in the network backbone as measured with the minimum spanning tree (MST). We found that TBI induced a shift in the MST toward weaker (MST: D = 0.08, p = 1.65 × 10^−7^) and lower cost connections (MST-Cost: D = 0.07, p = 2.3 × 10^−6^) (Figure 2D and E, respectively). Lastly, we assessed for an association between connections found in the NBS and MST. We found significant differences in the percentage of connections in common between the NBS and MST subnetworks between the two groups (T_17_ = 16.6, p = 5.8 × 10^−12^) (Figure 2F). Specifically, in the TBI group 12.3% ± 1.68 (SD) of connections were present in both subnetworks, while the Control group only 2.1% ± 0.98 (SD) were present in both subnetworks.

### Local Network Analysis

Our previous analysis identified a series of changes in subnetwork network structure, indicating that injury induced changes might be better assessed using local network measures. We therefore examined how network measures that assessed connectivity at individual brain regions could be used to measure local changes in network structure post-TBI. We first looked at three commonly used measures of regional connectivity that have previously been used to measure TBI-induced changes in brain structure: degree, betweenness-centrality, and local efficiency. As seen in Figure 3A-C, for each measure we were able to detect specific brain regions with increases/decreases in these measures relative to controls, but the spatial patterning of where these changes occurred differed between measures, suggesting that a multitude of different types of changes in local network structure are occurring throughout the brain.

We followed up these observations with a further assessment of local connectivity and investigated the changes that TBI induces in motif coherence, which measures how strongly a brain region participates in specific patterns of connections (Materials and Methods; Figure 3D) (Onnela et al., 2005). In the TBI group, 9.3% of brain regions (N = 14 brain regions from n = 8 rats) increased in motif coherence, and 6% of brain regions (N = 9 brain regions from n = 8 rats) decreased in motif coherence relative to the Control group (Figure 3E). These results confirm that in the TBI group, we see changes in network structure occurring at the local level.

### Detection of Subgroups Within Heterogeneous TBI Populations

One of the hallmarks of TBI is the diverse response in patients (Sidaros et al., 2008; Kinnunen et al., 2011; Ghajari et al., 2017) with a subset of patients and animals developing post-traumatic epilepsy (Frey, 2003; Kharatishvili et al., 2006). This diverse response implies the existence of subpopulations within the TBI group due to different trajectories in pathological progression. To test for the existence of subpopulations within the TBI group, we designed a two-step unsupervised clustering procedure based on the previously described network measures. Clustering based on individual network measures resulted in the detection of 2 clusters in the TBI group with the exception of Motif 4-199 which contained 3 clusters (individual columns of Figure 4A). Using these results to perform our final clustering step, we identified two subgroups in the TBI population, which we labeled T1 (n = 3 rats), and T2 (n = 5 rats) subgroup (Figure 4B). Notably, when the same analysis was performed on the Control group, we did not Identify any subpopulations (Supplemental Figure 2), suggesting that the observed heterogeneity within the TBI group is directly related to network reorganization post injury.

**Figure 4.**
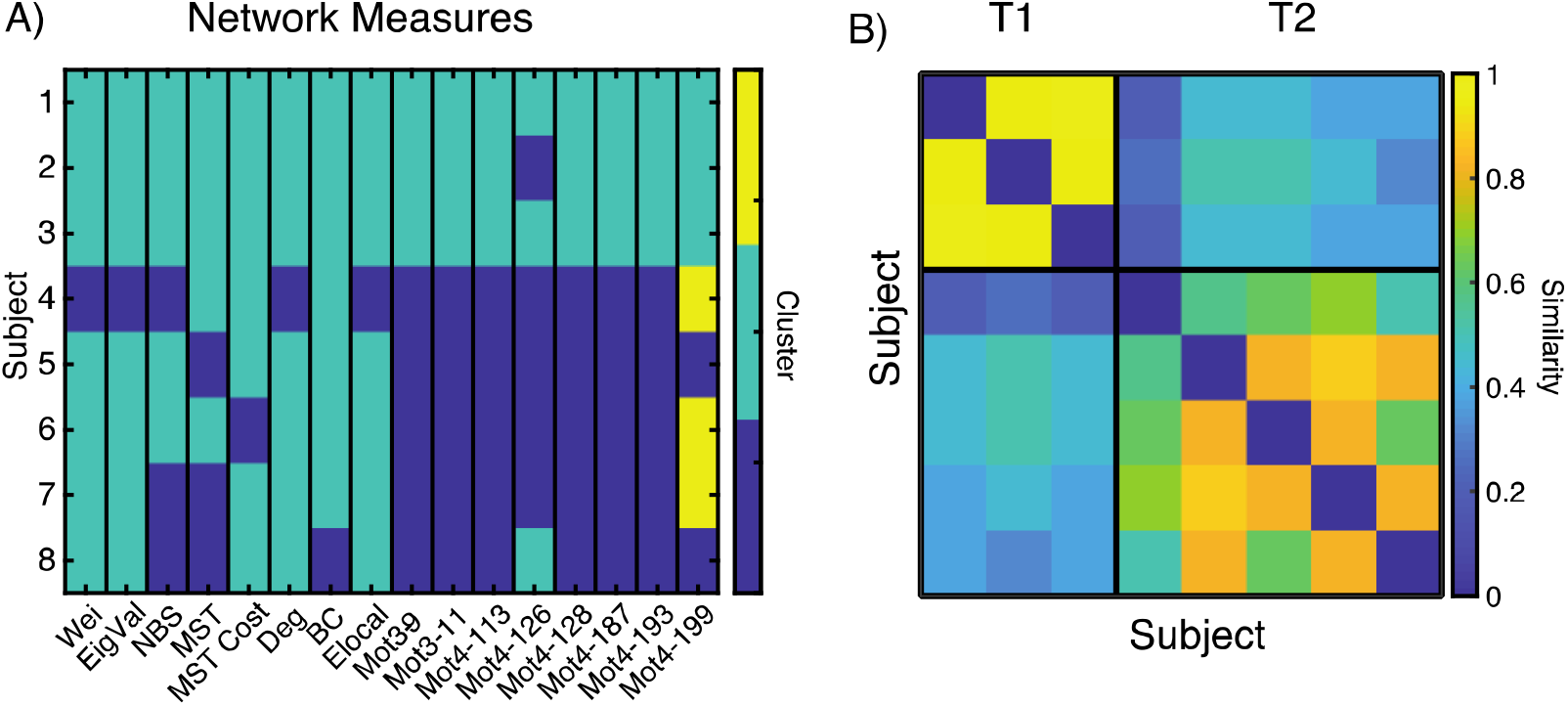
Clustering analysis and subpopulation identification of animals with TBI. A) For each network measure, *K-means* was applied to assign each animal to a single cluster. Columns represent the cluster identity of each animal clustered with a network measure. The color represents a unique cluster identity. B) Modularity-maximization clustering applied to the association matrix obtained from (A) was used to identify two subpopulations, T1 and T2, in the TBI group. The association matrix counts the fraction of times that two animals have the same cluster assignment in (A).

**Supplemental Figure 2.**
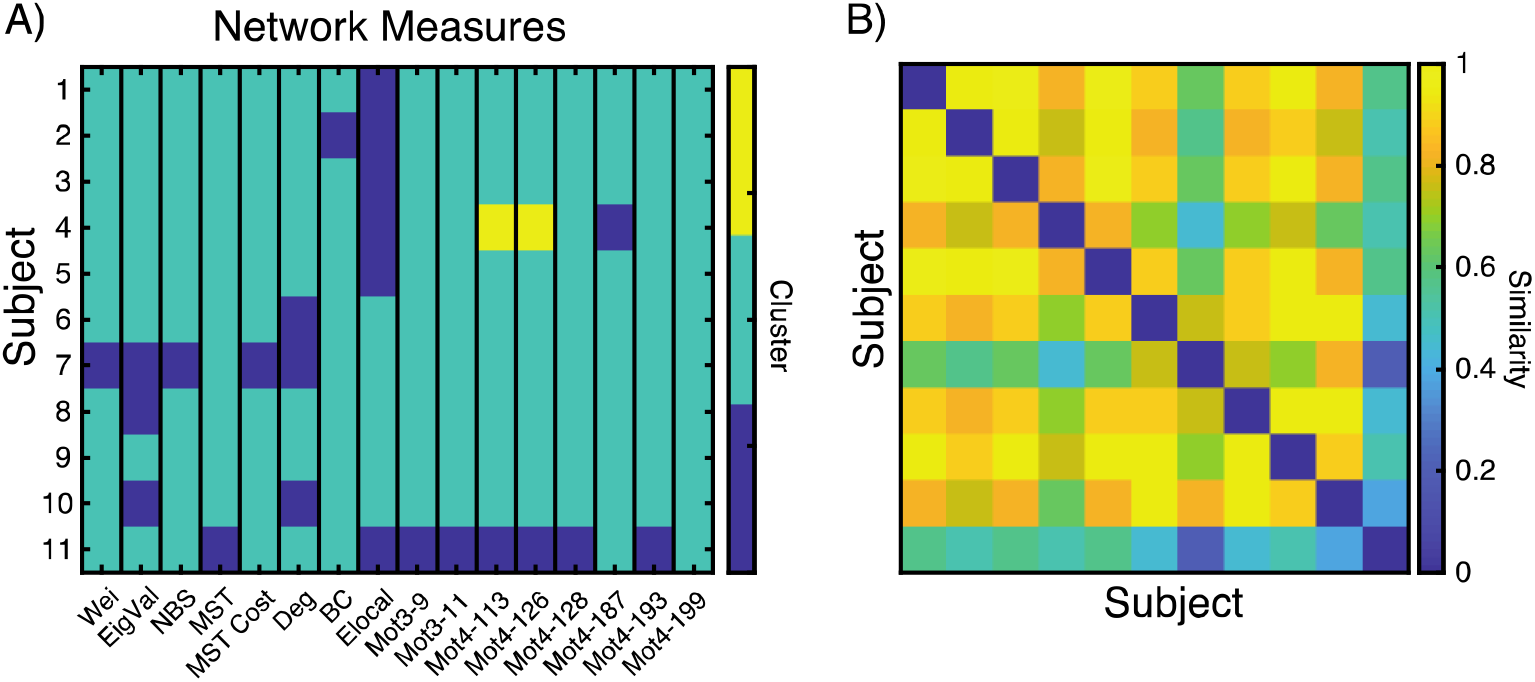
Clustering analysis of controls animals. A) For each network measure, *K-means* was applied to assign each animal to a single cluster. Columns represent the cluster identity of each animal clustered with a network measure. The color represents a unique cluster identity. B) Modularity-maximization clustering applied to the association matrix obtained from (A) determined that all animals were placed in the same group and no subgroups were present in the data.

### Network Motifs as Markers of Heterogeneity

When examining the membership of the T1 and T2 subgroups, we found that the T1 subgroup was composed of the first 3 animals, indicating that the motif analysis was driving the separation into two groups (see Figure 4A). To gain a better understanding of these changes, we analyzed how motif expression changed within the T1 and T2 subgroups relative to the control group. To account for the low number of animals, statistical testing was based on a permutation test in which subgroup labels were permutated 10,000x (see Materials and Methods). For the T1-subgroup, brain regions primarily increased in their motif coherence relative to Controls (Increase: N = 57 brain regions; Decrease: N = 1 brain regions from n = 3 rats) (Figure 5A). In contrast, in the T2-subgroup, brain regions primarily decreased in motif coherence (Decrease: N = 23 brain regions; Increase: N = 5 brain regions from n = 5 rats) (Figure 5B).

**Figure 5.**
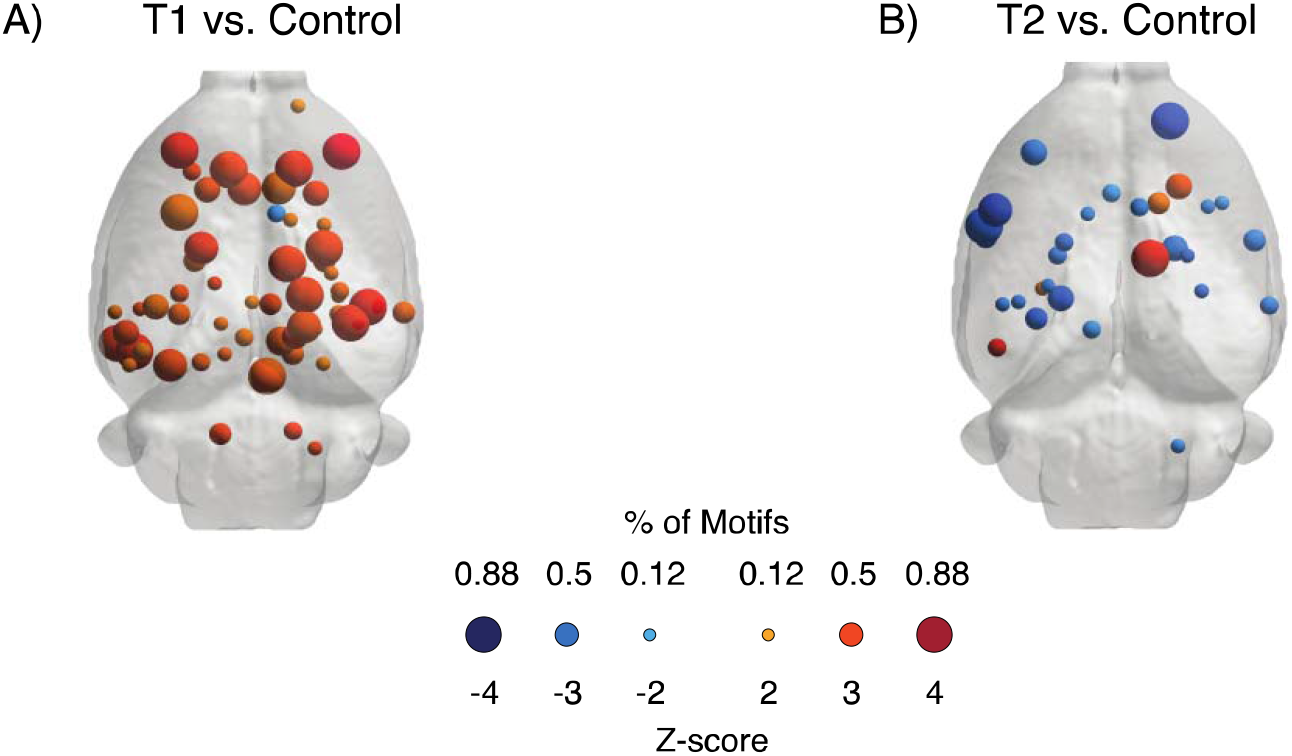
Subgroup analysis of regions for which differences in motif coherence are statistically significant. A) T1- and B) T2-subgroups for which a brain region’s motif coherence was significantly different from Controls. Color corresponds to the average Z-score of all significant motifs and size corresponds to percentage of motifs which a brain region was significantly different.

### Predicted Functional Connectivity

Given that we observed two subgroups with different types of structural changes in TBI animals, we next asked if these structural differences might lead to corresponding changes in functional brain activity. Previous work has indicated that the presence of different types of motifs can differentially impact the network’s propensity to promote synchronous brain activity (Terry et al., 2012). Functional connectivity describes statistical relationships between the activity of two brain regions, and the degree of functional connectivity can be associated with synchronization in brain activity. To assess potential differences in synchronized activity between the two subgroups, we tested if the structural differences observed between the T1- and T2-subgroups might result in the two subgroups displaying different patterns of functional connectivity. To predict functional activity of the two subgroups, we used a computational approach based on path transitivity (Goni et al., 2014) (see Materials and Methods).

In Figure 6A, we show the average predicted functional connectivity for the Control group, T1, and T2 subgroups. We found that the average change in functional connectivity was significantly increased for TBI group as a whole [TBI-Control: 0.008 ± 0.04 (SD)] and the T1-subgroup [T1-Control: 0.035 ± 0.06 (SD)], but decreased for and T2-subgroup [T2-Control: - 0.008 ± 0.05 (SD)] relative to Controls (F_2,32631_ = 1758, p << 0.0001) (Figure 6B).

**Figure 6.**
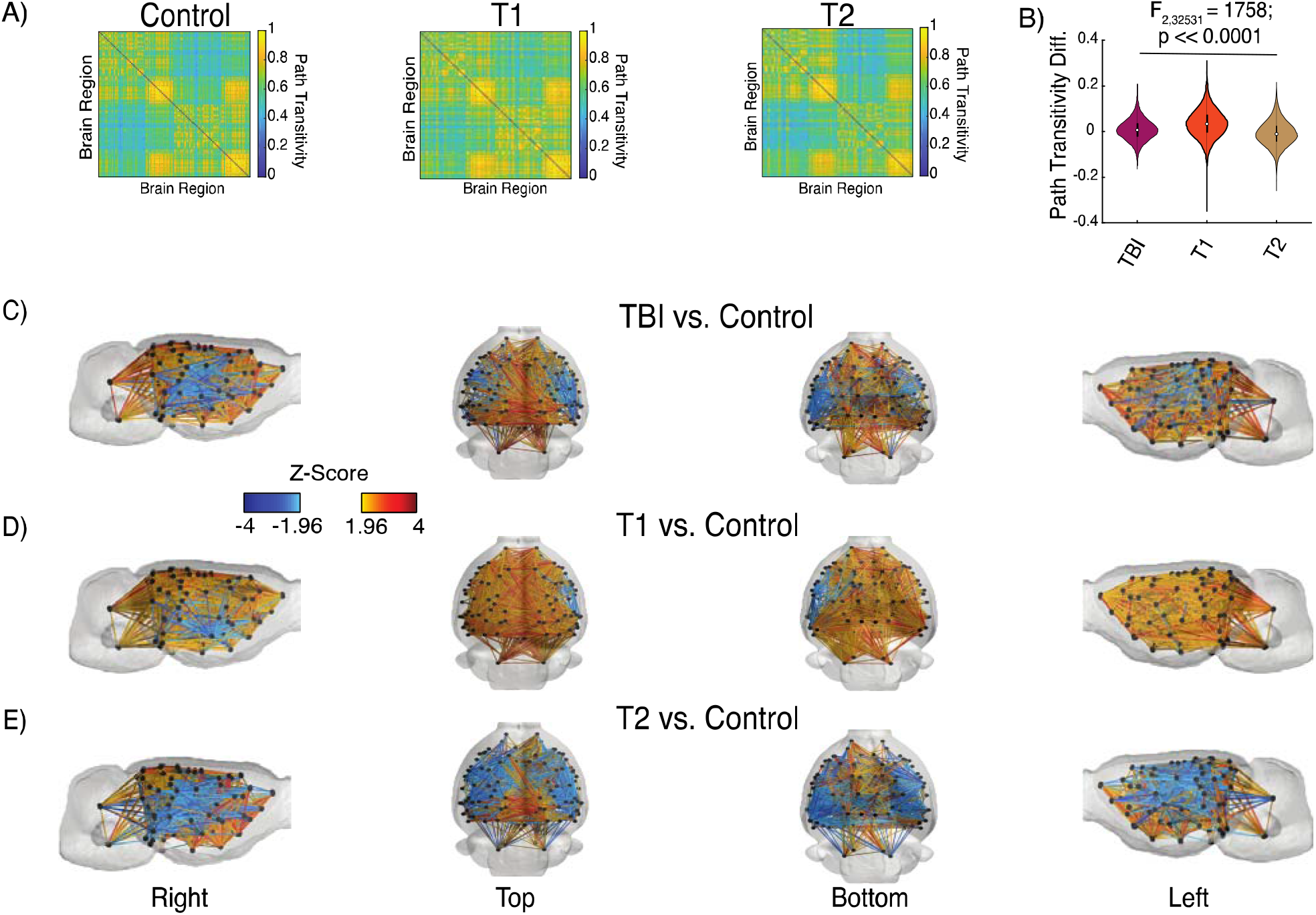
Predicted and changes in functional connectivity between TBI subgroups and Control group. A) Average predicted functional connectivity for the Control Group (Left), T1- (Middle) and T2- (Right) subgroups. B) Changes in average predicted functional connectivity (path transitivity) between TBI, T1- and T2-subgroups relative to the Control group. C-E) Individual connections which were significantly different in the TBI group, T1- and T2-subgroup from the Control groups, respectively.

Examining individual connections in predicted functional connectivity, we found that when compared to the Control group, in the TBI group 5.6% of connections increased in strength (N = 630 edges in n = 8 rats), while 1.9% of connections decreased in strength (N = 207 edges in n = 8 rats) (Figure 6C). Separating the animals into their respective subgroups, the T1-subgroup showed 9.2% of connections increased (N = 1033 edges in n = 3 rats) and 0.7% of connections decreased in strength (N = 81 edges in n = 3 rats) (Figure 6D). In contrast, the T2-subgroup showed 3.8% of connections increased (N = 422 edges in n = 5 rats) and 4.3% of connections decreased in strength (N = 483 edges in n = 5 rats) (Figure 6E). Interestingly, in both the T1- and T2-subgroups the connections that decreased in strength were located adjacent to the site of the injury.

## Discussion

TBI is marked by focal and diffuse jury to both grey and white matter in the brain (Gentleman et al., 1995; Smith et al., 2003; Johnson et al., 2013b). Along with structural damage, TBI has been associated with a variety of adverse functional/behavioral changes. For example, TBI increases the risk of developing post-traumatic epilepsy (Annegers et al., 1998; Englander et al., 2003; Christensen et al., 2009; Wang et al., 2013; Mahler et al., 2015; Webb et al., 2015; Smith et al., 2018) and it has been shown that a subpopulation of TBI patients with a similar pattern of abnormal network connections exhibited decreased cognitive performance (Solmaz et al., 2017). However, no clear biomarkers exist for predicting who will develop adverse functional effects, so it is important to characterize the heterogeneity of brain structural changes resulting from injury (Pitkänen and Immonen, 2014). Here we use network theory to characterize changes in global and local connectivity following TBI in rats induced by FPI in order to probe how heterogeneity in structural changes might drive some of the heterogeneity observed functionally and behaviorally.

We found that two global measures of structural connectivity were not able to distinguish the TBI group from the Control group, despite animals in the TBI group receiving a severe injury, which was validated by functional assessment (NSS). Instead we found changes in a subset of connections and nodes which is consistent with previous work that found changes in the density of a subset of fibers in moderate-to-severe injuries (Harris et al., 2016a; Iraji et al., 2016; Wright et al., 2017), and in subnetworks in structural connectivity following TBI (Hayes et al., 2016; Iraji et al., 2016; Thengone et al., 2016; Dall’Acqua et al., 2017).

These alterations indicate adaptive and maladaptive changes in brain connectivity (Fornito et al., 2015). Adaptive responses could indicate myelin repair (Armstrong et al., 2016) and/or axonal sprouting post TBI (Christman et al., 1997; Harris et al., 2010). On the other hand, maladaptive changes could be a result of axonal death (Gentleman et al., 1995; Smith et al., 2003), myelin damage (Bramlett and Dietrich, 2002; Johnson et al., 2013b) and neuroinflammation (Johnson et al., 2013a).

The changes in local connectivity were primarily found in shorter distance connections which increased in strength causing the network backbone to include more short-range connections. These shifts in the network backbone could be indicative of a break down in efficient communication between brain regions (Avena-Koenigsberger et al., 2017). In humans, moderate to severe TBI has been found to induce subject-specific alterations in the underlying white matter raising the possibility of individualized pathological trajectories (Ware et al., 2017). In our analysis, changes in brain structure could separate animals in the TBI group into two subpopulations, which we called the T1- and T2-subgroups. This aligns with previous work in humans in which alteration in structural connectivity in a population with moderate-to-severe injury separated individuals into subpopulations with corresponding differences in cognitive performance (Solmaz et al., 2017).

In our analysis, we found that the presence of subgroups was driven by differences in motif participation, indicating that local connections between brain regions could play an important role in network reorganization post-injury. Modeling studies that have focused on the relationship between motifs and synchronized activity have found that certain structural connectivity patterns, linear connections, Motif 3-9 (Figure 3D), promote zero-phase lag synchrony (Gollo and Breakspear, 2014). At the level of neuronal connectivity, computational modeling studies have shown that the presence of certain motifs relates to sustained neural activity (Bojanek et al., 2020). Additionally, recent modeling work identified the presence of superhubs – highly connected neurons that drive network activity through feedforward motifs – in epileptic networks (Hadjiabadi et al., 2021). Thus, the reorganization and increase of motif coherence in the T1 subgroup suggests that this subgroup might be expected to experience a change in brain activity, such as synchronization, as well.

The relationship between an increase in motif coherence and synchronization is supported by our predicted functional connectivity analysis in which we found that the T1 subgroup had an in increase in predicted functional connectivity relative to both the Control group and the T2 subgroup. This finding corroborates previous analyses that have found that severe TBI increases functional connectivity in humans (Mayer et al., 2011; Sharp et al., 2011; Hillary et al., 2014) and rats (Harris et al., 2016b). This is important given that individuals that sustain a severe TBI are at a 17 times greater risk of developing epilepsy (Annengers et al., 1998; Banerjee et al., 2009; Houser et al., 1990), and simulation models of epilepsy have found that focal, secondary- and primary-generalized seizure can arise from changes in network architecture (Terry et al., 2012). We might speculate that this subgroup could represent animals that are more likely to go on to develop PTE. However, the animals were not monitored using video-EEG to assess long term outcomes so future work would be needed to test this hypothesis. Despite this limitation, our analysis shows that network measures can be used to identify meaningful differences within a population and offers an insight pertaining to the disease progression for an individual.

In our clustering analysis, we were able to identify two distinct subgroups within the TBI group. While this grouping was driven by motif participation, one can see when examining the columns of Figure 3A that the structural connectivity (Wei), eigenvalue spectrum (EigVal), degree (Deg), and local efficiency (Elocal) result in the same groupings of animals, grouping all animals in the same cluster, with the exception of animal 4. This suggests that a hierarchical structure might also exist in the population, with sub-subgroups existing within subgroups. While the current data set is too small to fully examine such possibilities, this will be important to examine in future work with larger populations.

While our work highlights the importance of taking into account population heterogeneity when analyzing brain network data, it also contains several limitations. In this study, in each animal, the same injury was delivered to the same location. Despite this fact, we perhaps surprisingly still see heterogeneity in how the network reorganized post-injury. While in some ways, this is a strength of the study because it allows us to examine heterogeneity in a controlled environment, is also raises a question of if the observed changes in structural networks are occurring because this specific type of injury tends to produce certain changes in network structure. This is especially important because human TBI populations have injuries that are more diverse in severity and location. Despite this challenge, work in humans has found similar changes in a subset of connections in both moderate (Thengone et al., 2016; Dall’Acqua et al., 2017) and moderate-to-severe TBI populations (Solmaz et al., 2017). Additionally, subpopulations in human with moderate-to-severe TBI have been found based on changes in strucutral connectivity (Solmaz et al., 2017). Our analysis broadly reflects what has been found in the human TBI population, but future studies that incorporate different levels of severity and different in injury locations are required to translate our results to humans.

Another limitation of this work pertains to the relationship between structural and functional networks. As we do not have functional imaging data from which to calculate functional connectivity, in line with previous computational work (Honey and Sporns, 2008; Alstott et al., 2009; Olmi et al., 2019), we instead took a computational approach and simulated functional connections using the observed structural connectivity. The simulated functional connectivity is derived using path transitivity of the structural connectivity and only approximates observed patterns of resting state functional connectivity. However, the fact that we observed significant differences in simulated functional connectivity between the T1 and T2 subgroups relative to the Control group remains important as it is not necessarily the case that all changes in network structure would lead to significant changes in path transitivity. This inspires further work to more closely examine heterogeneity in experimentally derived functional connectivity derived from TBI populations.

Because TBI is a complex and heterogeneous disorder, where similar insults can give rise to different behavioral outcomes, it is essential to develop new analytical approaches that are tailored to the heterogeneities present within the data. Most work that searches for biomarkers of PTE looks for correlations between a single observed variable and the development of PTE. However, our findings suggest that future work should examine and embrace heterogeneity within populations to more effectively link specific patterns of brain reorganization to functional outcomes.

## Author Contributions

J.N. designed the analysis, processed the data, interpreted the data, and wrote/edited the manuscript.

M.G. designed the study, acquired data.

F.S. processed the data, acquired data, and edited the manuscript.

D.P designed the study, acquired data, and edited the manuscript.

S.F.M. designed the analysis, interpreted the data, and wrote/edited the manuscript.

